# Networks of common inputs to motor neurons of the lower limb reveal neural synergies that only partly overlap with muscle innervation

**DOI:** 10.1101/2021.10.13.460524

**Authors:** François Hug, Simon Avrillon, Aurélie Sarcher, Alessandro Del Vecchio, Dario Farina

## Abstract

Movements are reportedly controlled through the combination of synergies that generate specific motor outputs by imposing an activation pattern on a group of muscles. To date, the smallest unit of analysis has been the muscle. In this human study, we decoded the spiking activities of spinal motor neurons innervating six lower limb muscles during an isometric multi-joint task. We identified their common low-frequency components, from which networks of common synaptic inputs to the motor neurons were derived. The vast majority of the identified motor neurons shared common inputs with other motor neuron(s). In addition, groups of motor neurons were partly decoupled from their innervated muscle, such that motor neurons innervating the same muscle did not necessarily receive common inputs. Conversely, some motor neurons from different muscles – including distant muscles – received common inputs. Our results provide evidence of a synergistic control of a multi-joint motor task at the spinal motor-neuron level.

**Teaser:** The generation of movement involves the activation of many spinal motor neurons from multiple muscles. A central and unresolved question is how these motor neurons are controlled to allow flexibility for adaptation to various mechanical constraints. Since the computational load of controlling each motor neuron independently would be extremely large, the central nervous system presumably adopts dimensionality reduction. We identified networks of functional connectivity between spinal motor neurons based on the common synaptic inputs they receive during a multi-joint task. Our findings revealed functional groupings of motor neurons in a low dimensional space. These groups did not necessarily overlap with the muscle anatomy. We provide a new neural framework for a deeper understanding of movement control in health and disease.

## Introduction

It has been proposed that movements are controlled through a combination of muscle synergies. Each synergy is a functional unit that generates a motor output by imposing a specific activation pattern on a group of muscles (*1, 2*). By significantly reducing the number of controlled dimensions, the abovementioned control strategy is thought to simplify the production of movement.

Current approaches to assess muscle synergies rely on the rationale that the entire motor neuron pool innervating a muscle receives the same inputs. However, the concept of common inputs shared by entire pools of motor neurons has been challenged by recent studies. For example, human motor unit studies have identified motor unit behaviors that do not reflect the presence of common inputs shared by a motor neuron pool (i.e. innervating a single muscle) during finger flexion tasks (*3*) and multi-muscle grasping tasks (*4*). Moreover, in non-human primates, Marshall et al. (*5*) recently identified neural substrates that would allow independent control even at the single motor neuron level within a motor neuron pool. Further, they postulated that a common input shared by motor neurons is not a fundamental feature of the neural control of movement (*5*). Rather, movement may be more flexibly organized even at the level of an independent control of single motor units within a single muscle. However, such independent control of even a small proportion of motor units would imply a very large dimensional control space, orders of magnitude greater than the space determined by the number of muscles active in a given task.

While an independent control of single motor units may be possible, an observation of differential control between individual motor units does not necessarily imply that they could be controlled independently of all other motor units. In addition, different motor units, even within the same muscle, may belong to different functional groups receiving different common inputs (*3*). The control of motor units individually is unlikely to be justified by functional benefits. Indeed, the effect of independent inputs on force generation is negligible as force modulation is mainly influenced by the common synaptic input received by a population of motor neurons (*6, 7*). Accordingly, Bracklein et al. (*8*) observed that it is very challenging for humans to voluntarily disrupt the common input to motor neurons innervating a single muscle and thus to achieve independent control. There is a greater likelihood that movement is controlled via common inputs to groups of motor neurons than via an independent control of individual motor neurons. These groups of motor neurons may be partly decoupled from the innervated muscles, such that motor neurons innervating the same muscle may not necessarily receive the same inputs, while motor neurons from different muscles may receive the same inputs. This would imply a partial mismatch between muscle anatomy and the distribution of common inputs to the innervating motor neurons. Notably, the conventional approach of using electromyography (EMG) amplitude recorded from many muscles to identify muscle synergies (*1, 2*) does not allow for the distinction between the possibility of independent or common control at the level of groups of motor neurons within and across muscles.

In this study, we identified groups of motor neurons receiving common inputs during an isometric multi-joint task. Our analysis was performed using a unique dataset of dozens of spinal motor neurons from six lower limb muscles. We did not impose any *a priori* muscle anatomical constraints in the identification of common inputs to motor neurons; however, we used a purely data-driven method to identify the groups of motor neurons based on their level of common low-frequency modulation in discharge rate. With this approach, we built networks of common inputs to motor neurons based on natural motor neuron behavior. We hypothesized that each motor neuron would group with others in functional clusters, based on the common input received with other motor neurons. Furthermore, we hypothesized that an independent modulation of single motor neurons would be very rarely, if at all, observed. If these hypotheses are supported, the observations would provide evidence that a common input is an essential feature of the neural control of movement at the motor neuron level. As secondary hypothesis, we predicted that motor neuron grouping solely based on common inputs would not necessarily correspond to the muscle innervation, but rather to functional associations. Accordingly, we expected that some motor neurons from distant pools could receive a common input, consistent with their combined role in endpoint force orientation.

## Results

### Identification of motor neuron activity during an isometric multi-joint task

Ten participants were instructed to perform a multi-joint task, which consisted of producing a submaximal force with their right leg on an instrumented pedal, while seated on a cyclo-ergometer (Fig. 1A). The crank was fixed at 135° from the top dead center such that the task was isometric. Using a real-time feedback, participants were asked to match both the pedal angle and the total reaction force vector corresponding to those produced during a submaximal dynamic cycling task at the same crank angle (see Methods). All the participants were able to match the targets, as demonstrated by the low root mean square error values of the pedal angle (2.0 ± 1.3°), and the total reaction force vector angle (1.1 ± 0.5°), as well as its norm (4.7 ± 0.9 N) (Fig. 1B).

**Fig 1.**
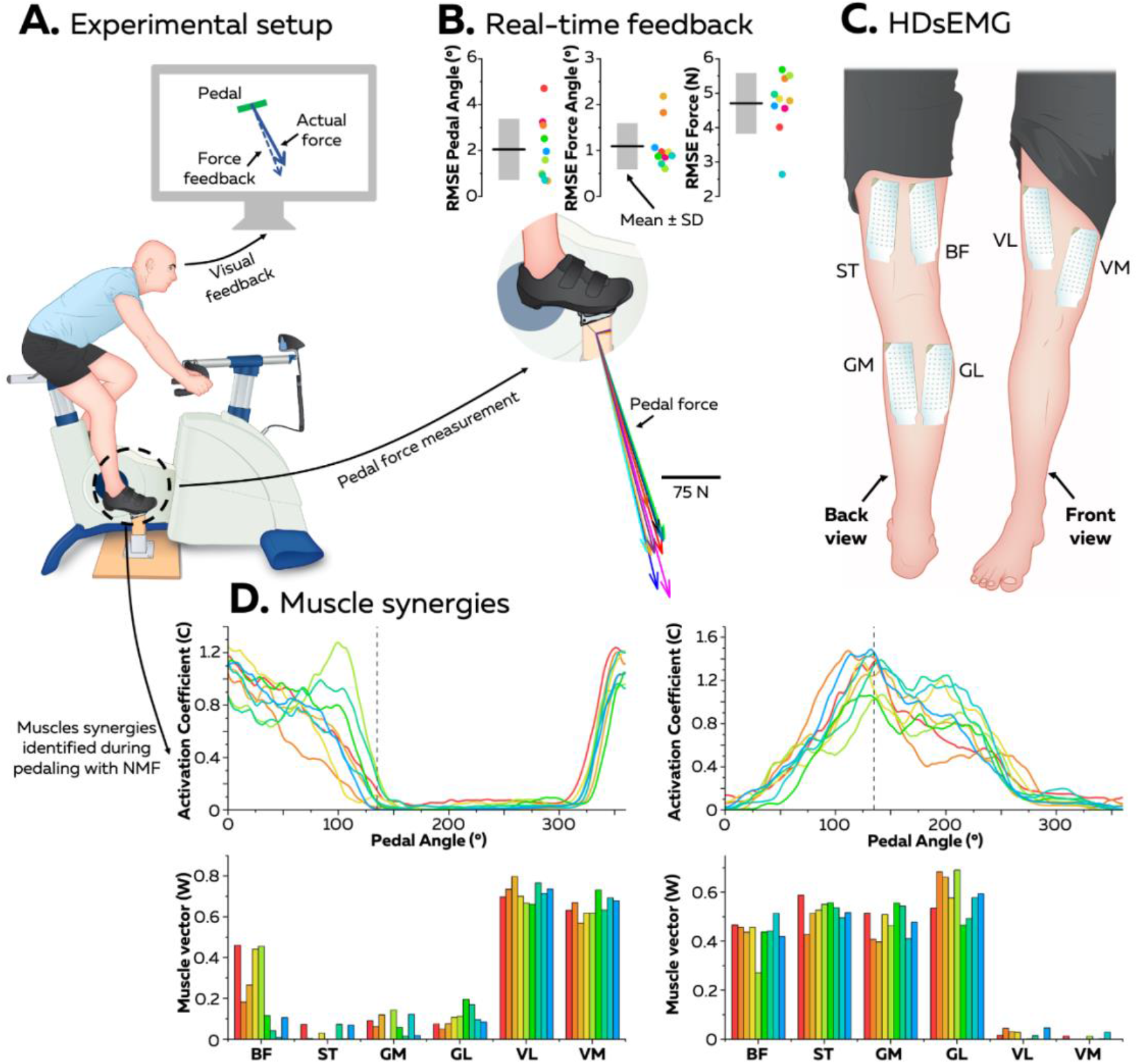
Experimental measures. An experimental session involved an isometric multi-joint task, wherein a participant matched a force vector on an instrumented clipless pedal while seated on a cyclo-ergometer. A real-time force feedback was provided to the participants (Panel A). The total reaction force vector produced by the participants is depicted in Panel B (each participant [n=10] is represented by a different color). The root mean squared error (RMSE) is reported for the pedal angle (°), the total reaction force vector angle (°), and its norm (N) (Panel B). During this task, high-density EMG signals (HDsEMG) were recorded with a grid of 64 electrodes from six lower limb muscles of the right leg (Panel C). Muscle synergies were also identified during a dynamic pedaling task using non-negative matrix factorization applied to conventional bipolar EMG signals. Two synergies were identified (Panel D – each color corresponds to a participant). Notably, at the crank angle chosen for the isometric task (135°, vertical dashed line on the activation coefficients), mainly the second synergy involving the gastrocnemii and the hamstrings was active. biceps femoris (long head, BF), semitendinosus (ST), gastrocnemius medialis (GM), gastrocnemius lateralis (GL), vastus lateralis (VL), and vastus medialis (VM), NMF, non-negative matrix factorization.

We recorded high-density surface EMG (HDsEMG) signals from six lower limb muscles during two contractions, each lasting approximately 40 s and interspaced by 10 s of rest (Fig. 1C): biceps femoris (long head, BF), semitendinosus (ST), gastrocnemius medialis (GM), gastrocnemius lateralis (GL), vastus lateralis (VL), and vastus medialis (VM). An algorithm was used to separate these signals into motor unit spike trains. The number of motor units considered for each analysis is reported in Table 1. Of note, because there is a one-to-one relationship between the generation of an action potential in the innervated muscle fibers and the generation of an action potential in the motor neuron, we refer to motor unit or motor neuron spike trains indifferently throughout the article.

**Table 1.** Number of motor units considered for each analysis

| Muscle | Repeatability analysis |  |  | Main analysis |  |  |
| --- | --- | --- | --- | --- | --- | --- |
| | Sum | Mean ± SD | Range | Sum | Mean ± SD | Range |
| **BF** | 53 | $7.6 \pm 3.1$ | 4-14 | 91 | $9.1 \pm 5.1$ | 2-21 |
| **ST** | 21 | $3.0 \pm 4.3$ | 0-12 | 54 | $5.4 \pm 4.1$ | 1-12 |
| **GM** | 111 | $15.9 \pm 6.3$ | 9-26 | 168 | $16.8 \pm 9.6$ | 3-36 |
| **GL** | 38 | $5.4 \pm 6.0$ | 0-15 | 53 | $5.3 \pm 7.4$ | 1-22 |
| **VL** | 58 | $8.3 \pm 4.3$ | 0-13 | 81 | $8.1 \pm 3.7$ | 3-14 |
| **VM** | 28 | $4.0 \pm 2.3$ | 0-7 | 42 | $4.2 \pm 2.6$ | 2-9 |

SD, standard deviation ; BF, biceps femoris; ST, semitendinosus; GM, gastrocnemius medialis; GL, gastrocnemius lateralis; VL, vastus lateralis; VM, vastus medialis.

### Common inputs to motor neurons

We estimated the common input shared between motor neurons by calculating the cross-correlation between smoothed discharge rates for each pair of motor units (Fig. 2B). The smoothed discharge rates were obtained by convoluting the binary signals (motor unit spike times) with a 400-ms Hanning window. The length of the Hanning window was chosen such that the correlation was calculated based on the low-frequency oscillations of the signal [2.5 Hz, effective neural drive (*6*)] thereby limiting the effect of the nonlinear relationship between the synaptic input and the output signal (*9*). Initially, we assessed the repeatability of the correlation matrices (Fig 2C) between the two contractions interspaced by 10 s of rest. Specifically, we selected two 10-s windows containing the activity of the same motor units. We were unable to match enough motor units between the two contractions for three participants; hence, this analysis was performed on the remaining seven participants, and therefore on a smaller number of motor units than for the main analysis described below. Specifically, a total of 309 motor units (mean per participant ± standard deviation: 44.1 ± 13.7) were considered in this repeatability analysis (Table 1).

**Fig 2.**
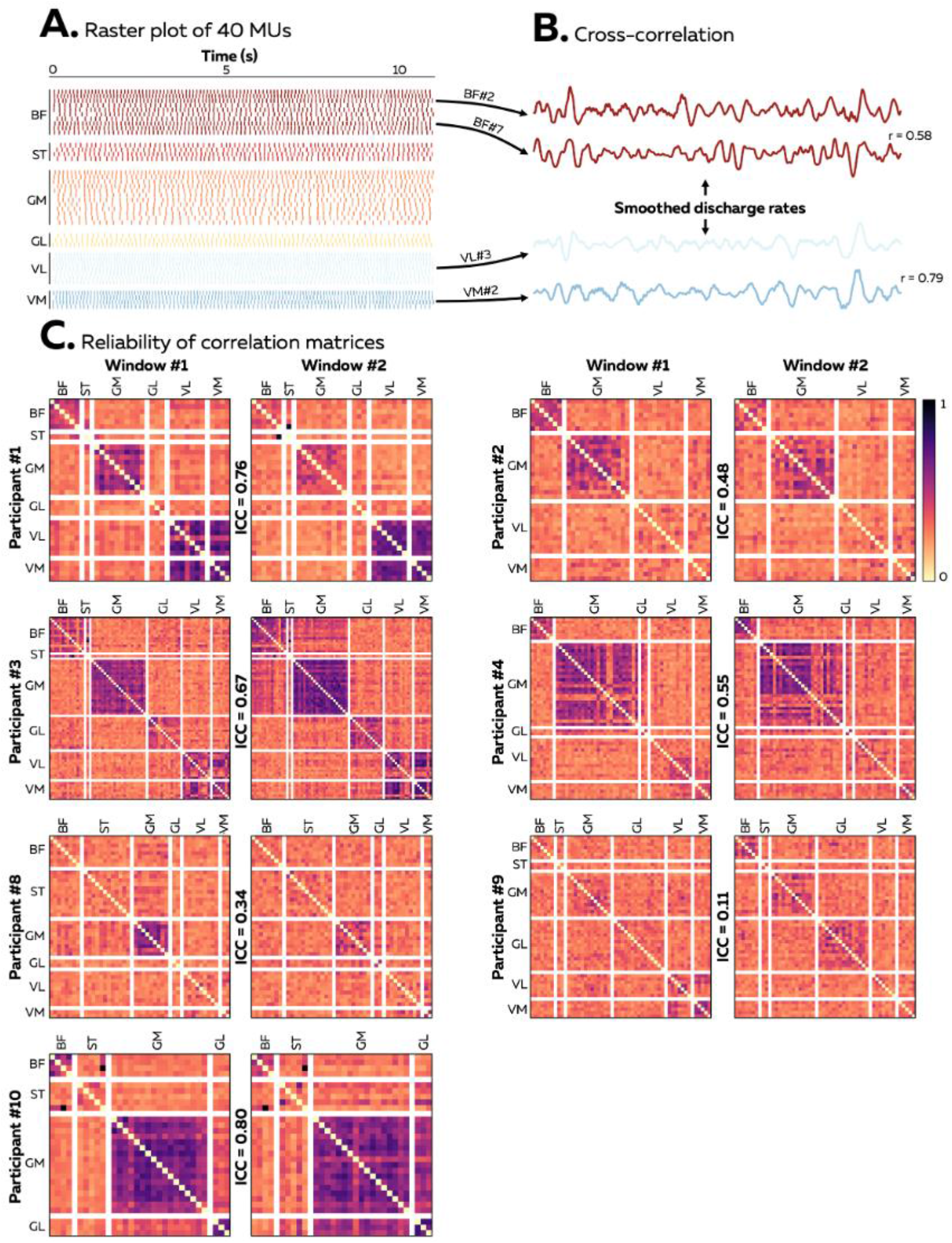
Within-session reliability of the correlation matrices. The decomposed motor unit discharge times were first converted into continuous binary signals (Panel A) and then convoluted with a 400-ms Hanning window to estimate the effective neural drive. A cross-correlation function was applied on the smoothed discharge rates (Panel B). Two 10-s windows with the same motor units were identified to assess the reliability of correlation matrices. We used Intraclass correlation coefficient (ICC) to assess the reliability of the correlation matrices that gather the cross-correlation coefficients for all the pairs of motor units (Panel C). The color scale displayed on the upper right panel indicates the strength of the correlation between 0 and 1. BF, biceps femoris; ST, semitendinosus; GM, gastrocnemius medialis; GL, gastrocnemius lateralis; VL, vastus lateralis; VM, vastus medialis.

The correlation matrices were built from pairwise cross-correlations performed on an average of 1032 (range: 406-1378) motor unit combinations per participant (Fig. 2C). This analysis revealed an overall moderate repeatability (mean intraclass correlation coefficient [ICC] = 0.53, range: 0.11 to 0.80), with some participants exhibiting an excellent repeatability (Fig. 2A). Of note, the lowest ICC valued found in participant #9 (0.11) could be explained by the overall weak correlation, and therefore to the lower variance within the matrix (Fig. 2C).

Overall, there was a moderate to low repeatability of the degree of correlation between motor neuron spiking activities, depending on the participant. This was presumably due to the dependance of the correlation measure to factors other than the common input to motor neurons, as previously described (*10*). Measures of correlation between spike trains cannot be used to quantify the exact correlation between inputs to motor neurons (*10, 11*), and therefore may show different levels of repeatability even when computed for the same pairs of motor neurons. To address this fundamental limitation, we proposed to threshold the correlation coefficients, based on statistical considerations, as described in the following section. We evaluated the repeatability of the outcome measures once we applied this approach and observed a substantial increase in reliability (as described in the following).

### Networks of common inputs to motor neurons

We used a purely data-driven approach grounded on graph theory to extract networks of motor neurons based on the common synaptic input (Fig. 3). To account for the fact that the strength of common synaptic inputs between two motor neurons does not necessarily translate into a proportional degree of correlation between their outputs (common drive) (*10*), the networks were based on the significance of the correlations between motor neuron spiking activities rather than on the strength of the correlations. Specifically, we used a thresholding approach wherein only significant correlations were considered to build the networks (Fig 3B). The threshold for significance was defined as the 99^th^ percentile of the correlation coefficient distribution generated with resampled versions of the motor unit spike trains (see Methods). This approach assumes that a significant correlation in output corresponds to a significant correlation in input (*7*). This assumption is based on the evidence that a total lack of a common input between two motor neurons inevitably determines independent spiking activities. Therefore, the networks reflect groups of motor neurons that share common synaptic inputs. The networks do not maintain any information on the strength of the correlation, which is highly variable depending on factors other than the strength of the common input.

**Fig 3.**
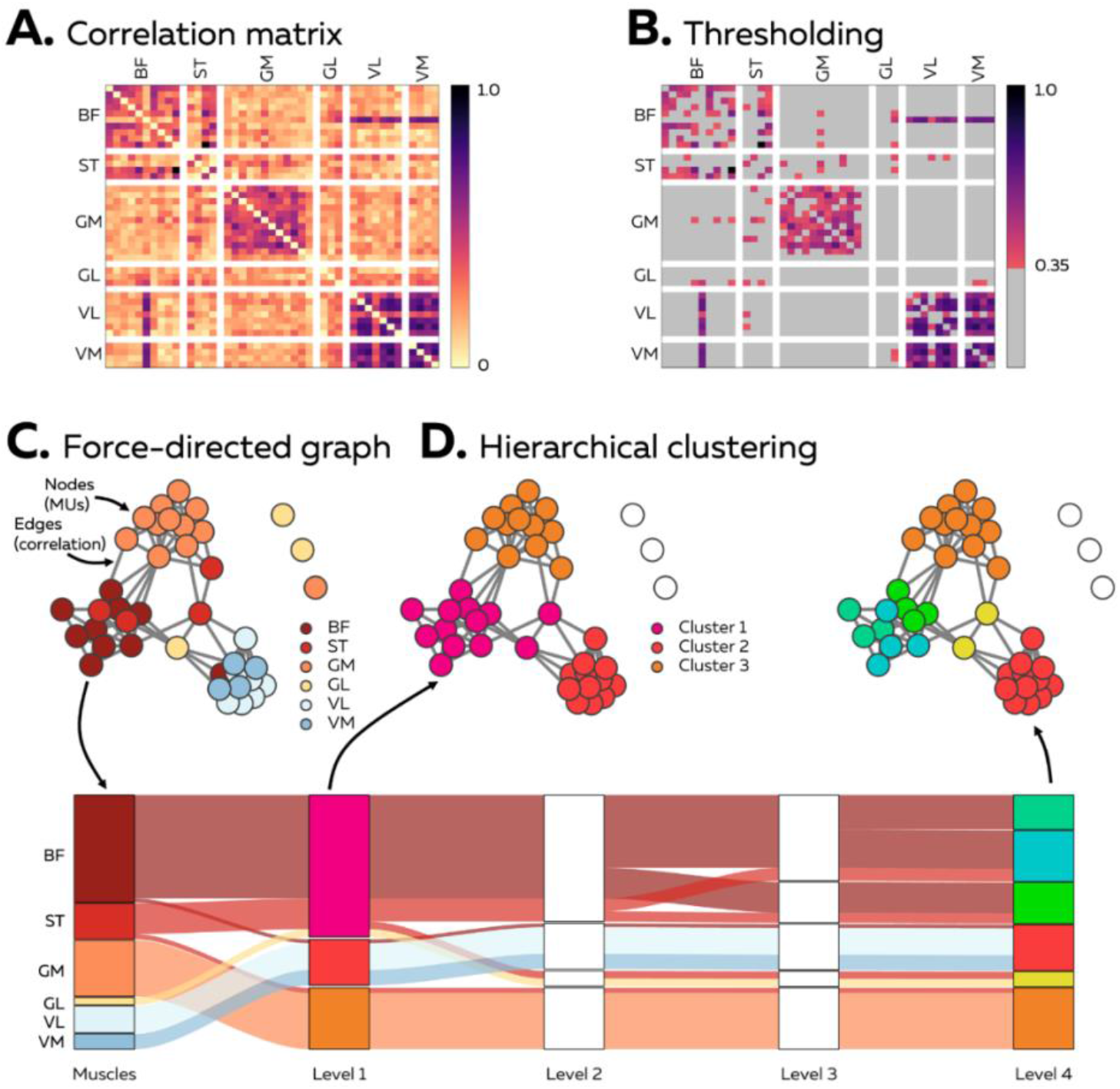
Construction of the networks of inputs to motor neurons. A cross-correlation function was applied on the smoothed discharge rates (Panel A); and only the coefficients of correlation that reached a significant threshold of 0.35 (see Methods) were considered to build the networks (non-gray squares in Panel B). Then, the networks were constructed using a force-directed graph with the Fruchterman-Reingold algorithm (Fruchterman & Reingold, 1991). Motor neurons (nodes) with a significant degree of common input, tend to attract each other, resulting in graphs having edges with uniform and short lengths, whereas nodes that are not connected tend to be positioned farther apart (Panel C). Finally, we applied a multiresolution consensus clustering approach to group the motor neurons according to their position and their connections in the graph (Panel D). This is an iterative procedure that converges toward a consensus matrix. This procedure stops once no additional statistically significant clusters can be identified (here, at level 4). Of note, the construction of the networks does not include any *a priori* information on motor neuron grouping (e.g., innervated muscles), and is therefore signal-based, without any anatomical constraint.

**Fig 4.**
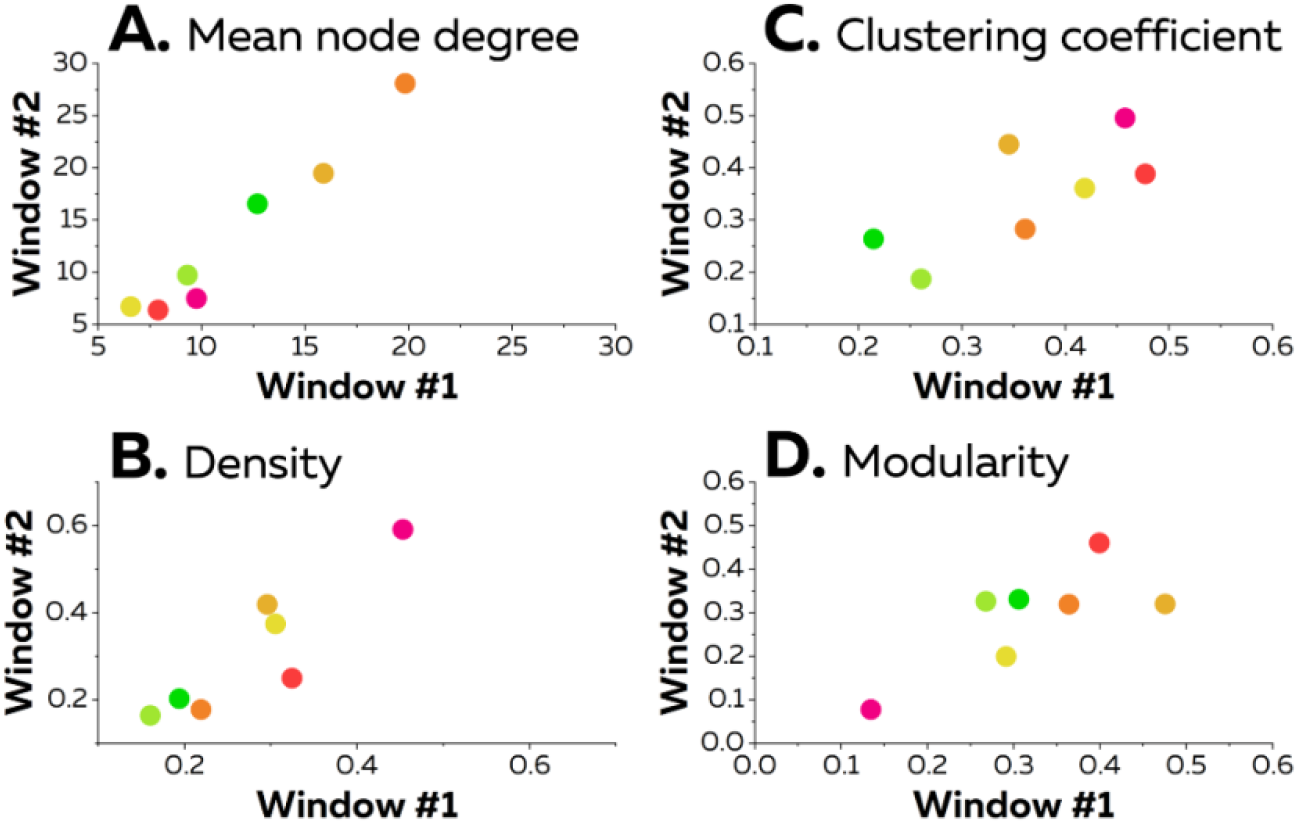
Within-session reliability of the networks. The reliability of four metrics that describe the properties of the networks was assessed between the two windows. Each scatter point represents an individual participant.

The networks were constructed using force-directed graphs with the Fruchterman-Reingold algorithm (Fig. 3C) (*12*). In these networks, motor neurons are nodes and edges are significant correlations between motor neurons. Nodes with strong connections - i.e., motor neurons with a significant degree of common input (correlation) - tend to attract each other, resulting in graphs having edges with uniform and short lengths, whereas nodes that are not connected tend to be positioned farther apart. The generation of the graph according to the above-described procedure does not include any a priori information on motor neuron grouping (e.g., innervated muscles), and is solely based on the significant correlations between motor neuron activities. Therefore, the procedure is purely signal-based, without any physiological or anatomical constraint. Finally, we applied a multiresolution consensus clustering approach to group the motor neurons according to their positions and their connections in the graph (Fig. 3D). This is an iterative procedure that stopped once no additional statistically significant clusters could be identified (consensus matrix).

First, the repeatability was assessed using classical network measures between the same two 10-s windows as those used to test the repeatability of the correlation matrices. The mean node degree, which corresponds to the mean number of edges connected to each node (i.e. the mean number of significant correlations for each motor neuron), exhibited a very good repeatability (ICC=0.85). Similarly, the density, which corresponds to the fraction of actual connections (significant correlations) to possible connections, exhibited a very good repeatability (ICC=0.81). Furthermore, the clustering coefficient, which is equivalent to the fraction of node neighbors that are neighbors of each other, exhibited a good repeatability (ICC=0.72). Finally, we observed a good repeatability for the modularity (ICC=0.75), which relies on the strength of division of the networks into communities (or clusters). Therefore, despite the values of correlation between motor neuron spike trains showed moderate repeatability, following thresholding, the network characteristics demonstrated good to very good repeatability.

Second, the main analysis was performed on one 10-s window selected such that the number of identified motor units was maximized. Consequently, this analysis was performed on a higher number of motor units than that used for the repeatability analysis. Specifically, a total of 489 motors units were considered, ranging from 20 to 99 per participant (Table 1). The mean discharge rates were 8.1 ± 1.0, 7.2 ± 1.1, 7.2 ± 1.0, 8.1 ± 1.5, 8.7 ± 1.4, and 9.0 ± 1.0 pps for BF, ST, GM, GL, VL, and VM, respectively.

As mentioned above, the motor neuron networks were constructed from the pairwise correlation coefficients that reached the significance threshold. The fraction of significant correlations to the total number of correlations (density) was 0.27 ± 0.12. Remarkably, an average of only 1.5 ± 1.5 motor neurons per participant (range: 0 to 4; 0 to 12% of the total number of motor neurons) were not connected to the network, i.e. they were not significantly correlated to any of the other motor neurons. This relatively small number of individual motor neurons that shared no common input with other identified motor neurons needs to be interpreted in relation to the limited sample of active motor neurons detected. There would likely be an even lower proportion of motor neurons without connections, if all the active motor neurons could be identified.

Fig. 5 depicts the correlation matrices and the networks for each participant. When considering the population level (all ten participants), four main observations can be made. First, motor neurons from the same muscles strongly overlap and are densely connected, highlighting a high level of within-muscle common input (Fig. 5 and 6). This was observed at a slightly lower extent for motor neurons from the ST muscle (Fig. 6). Second, VL and VM motor neurons overlapped and were densely connected in most of the participants meaning that common input was almost as high between motor neurons from the VL and VM pools as within the same pool (Fig. 5 and 6). This is further confirmed by the fact that some motor neurons from VL and VM were represented within the same cluster in 9 of 10 participants, with all of the motor neurons from VL and VM being in the same cluster in 6 participants (Fig. 7). Even though edges were observed between motor neurons from other anatomically defined synergist muscles (BF-ST and GM-GL), they were not as densely connected as those of VL-VM (Fig. 5). Indeed, some BF and ST, as well as GM and GL, motor neurons were represented within the same cluster in only 6 of 10 participants (Fig. 7). Third, edges were found between motor neurons from distant muscles, mostly between gastrocnemii and hamstring muscles. Specifically, some motor units from GM-BF, GM-ST, GL-BF, and GL-ST muscle pairs were represented within the same cluster in 6, 6, 3, and 4 participants, respectively. Notably, the local density between these muscles was relatively high in some participants (Fig. 6). Finally, it is noteworthy that some motor neurons from antagonist muscles, mostly BF-VL and BF-VM, were observed within the same cluster in 5 and 7 participants, respectively. However, the local density was relatively low; thus, only a very small sample of the motor neurons identified in these muscles were concerned.

**Fig. 5.**
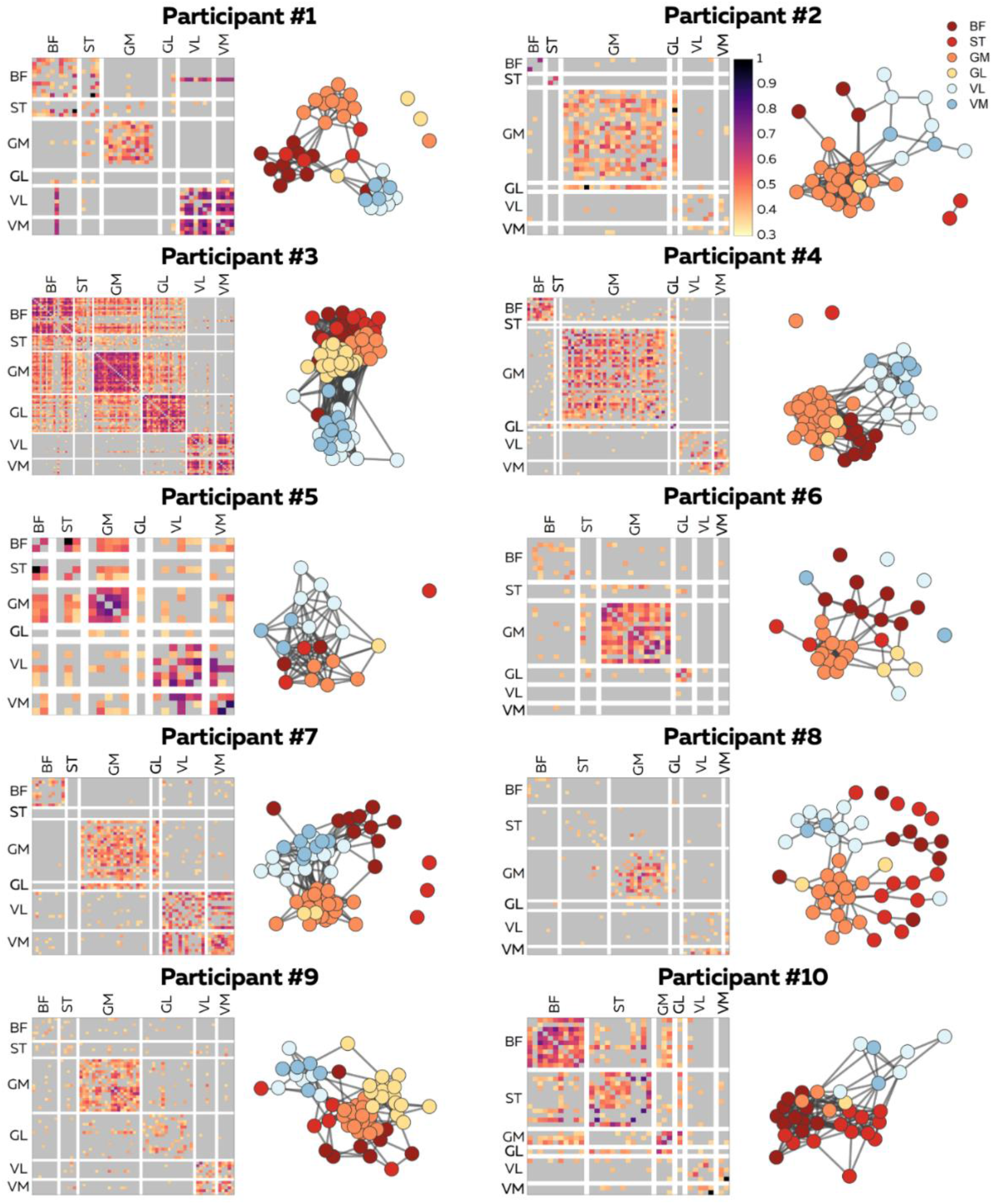
Networks of inputs to motor neurons. For each participant, a correlation matrix gathered all the correlations that exceeded a significant threshold of 0.35. The color map represents the strength of the correlation coefficient. However, this information is only presented here to facilitate the interpretation of the graphs, as we did not consider the strength of the correlation to build the networks. The networks were constructed using force-directed graphs. Each node represents a motor neuron and each edge represents a significant correlation between motor neurons, i.e., a significant common input to these motor neurons. BF, biceps femoris, ST, semitendinosus; GM, gastrocnemius medialis; GL, gastrocnemius lateralis; VL, vastus lateralis; VM, vastus medialis.

**Fig. 6.**
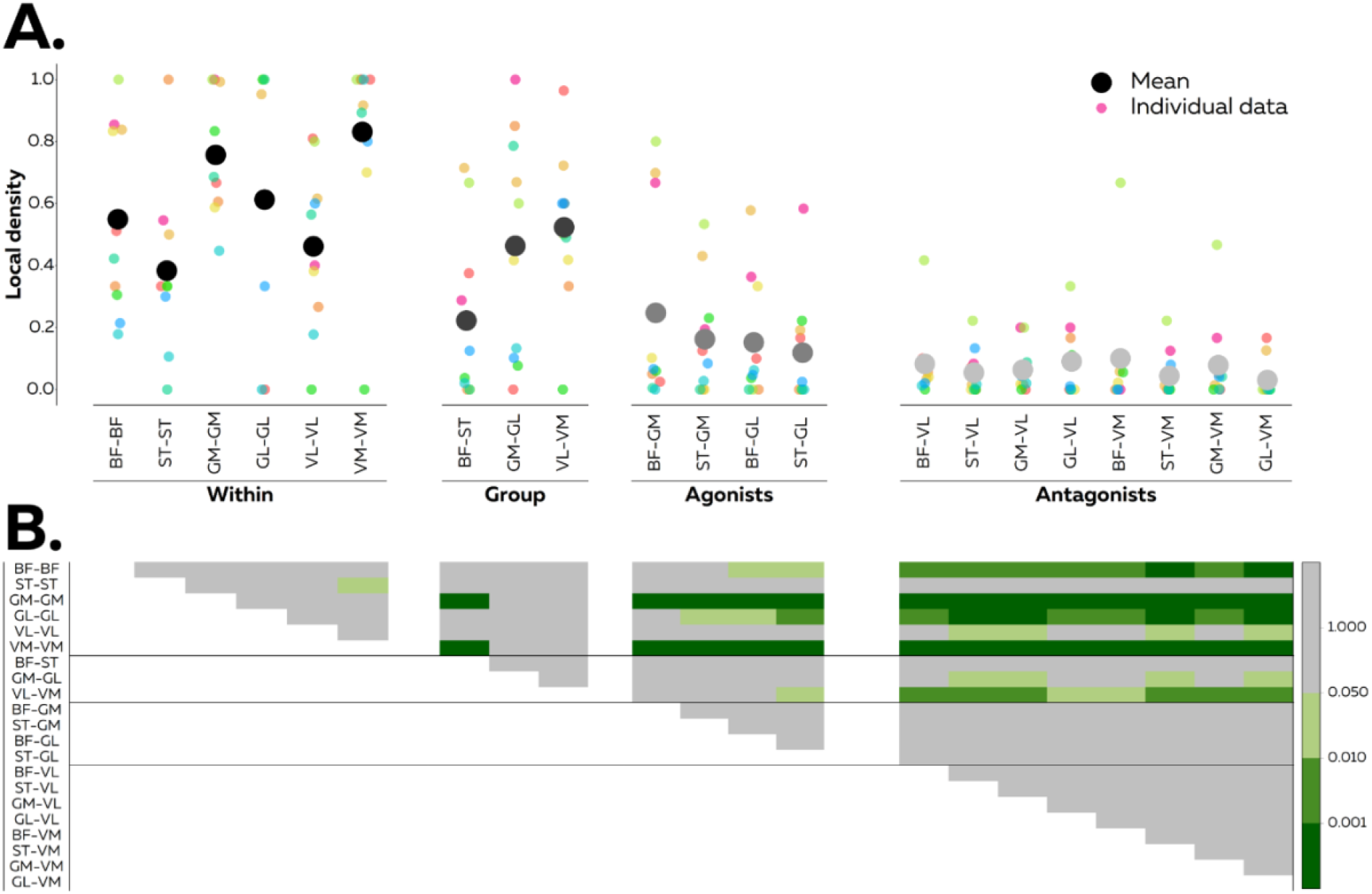
Local density for each muscle or muscle pair. The local density corresponds to ratio of the number of significant correlations to the total number of motor neuron pairs (Panel A). We performed this analysis on motor neurons from the same muscle (within) and on motor neurons from two different muscles which: i) belong to the same anatomical group (i.e., hamstrings, triceps surae, quadriceps), ii) share a common function (i.e., agonists), or iii) exhibit opposite functions (antagonists). Panel B displays the level of statistical significance of the multiple comparisons (one-way analysis of variance was applied on the local density with Bonferroni as post-hoc test). BF, biceps femoris, ST, semitendinosus; GM, gastrocnemius medialis; GL, gastrocnemius lateralis; VL, vastus lateralis; VM, vastus medialis.

**Fig 7.**
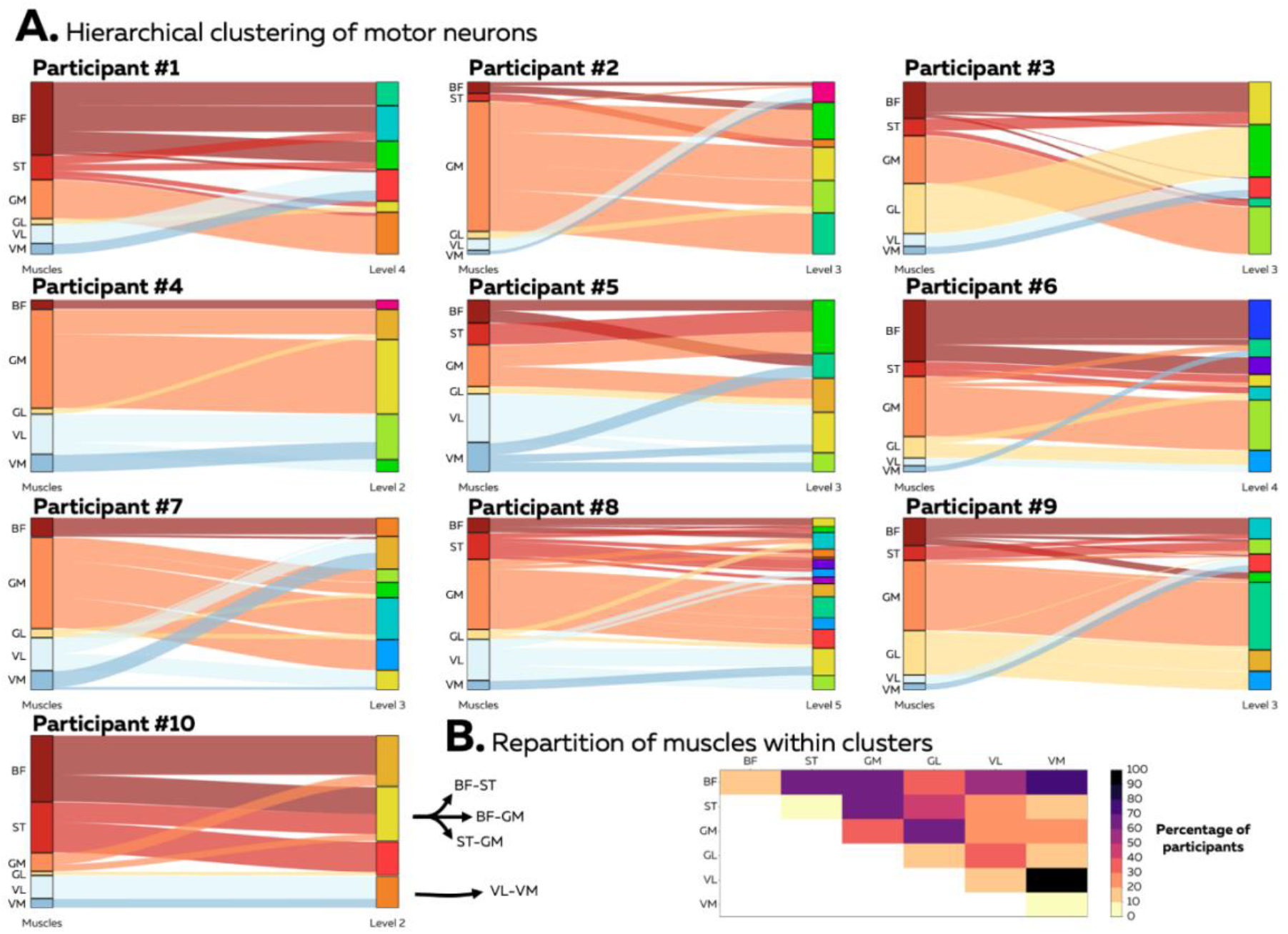
Clusters of motor neurons for each network. We applied a clustering procedure to group the motor neurons according to their positions in the graph, and therefore based on their common input. We defined a cluster as a group of motor neurons densely connected to each other and loosely connected to the rest of the network. We used a multiresolution consensus clustering method to identify significant clusters at different resolutions, i.e., levels (Panel A). As the clusters were decoupled from the muscle innervation, we also reported the occurrence of each muscle pair within the same cluster (Panel B). Cells with the same muscles in the x- and y-axes (e.g., BF-BF) represent the percentage of participants with a cluster that only groups motor neurons from this muscle. BF, biceps femoris, ST, semitendinosus; GM, gastrocnemius medialis; GL, gastrocnemius lateralis; VL, vastus lateralis; VM, vastus medialis.

### Reduced dimensionality

From the 47.4 ± 22.8 (range: 19-99) motor units per participant used to construct the networks, we identified 6.6±2.8 clusters from the consensus partition (see Methods). Of note, only five clusters were identified in participant #3 who exhibited the highest number of motor units (n=99). Overall, these results suggest that the control of the studied isometric multi-joint task is achieved through a large dimensionality reduction at the motor neuron level. Although the average number of clusters was comparable to the number of muscles (n=6), clusters were not anatomically defined, as the vast majority of them were composed by motor neurons from different muscles; moreover, motor neurons from different muscles could be represented in multiple clusters. Specifically, on average, each cluster was composed of motor neurons innervating averagely 1.9 ± 0.3 muscles (range across participants: 1.4-2.4). In addition, motor neurons from BF, ST, GM, GL, VL, and VM were represented in 2.7 ± 1.1, 2.2 ± 2.1, 2.9 ± 1.7, 1.6 ± 1.0, 1.8 ± 1.1, and 1.3 ± 0.7 clusters, respectively.

### Muscle synergy analysis

After the completion of the main protocol, HDsEMG electrodes were replaced with conventional bipolar electrodes and muscle synergies were identified using non-negative matrix factorization applied to the EMG signals recorded during a dynamic pedaling task. Two muscle synergies were identified (variance accounted for: 91.0 ± 2.5%; Fig 1D). At 135°, which corresponds to the angle used for the isometric task, only the second synergy composed by BF, ST, GM, and GL was active.

## Discussion

We used a purely data-driven method grounded on graph theory to extract motor neuron networks based on common synaptic inputs. The vast majority of the identified motor neurons shared common inputs with other motor neuron(s), with a partial mismatch between this distribution of common inputs to motor neurons and innervated muscles. The results provide evidence of a neural synergistic control of a multi-joint motor task at the spinal motor neuron level.

### Common input at the spinal motor neuron level

The concept of synergistic control of movement has received considerable attention since its inception by Bernstein (*13*). To date, the smallest unit of analysis has been the muscle through muscle activation assessment. This level of analysis inherently constraints the dimensionality of the neural control to be less than or equal to the number of recorded muscles. Even though this approach led to important advances in our understanding of the modularity of movement control in health (*14*) and disease (*15*), the muscle is not the lowest level of movement control. Rather, the spinal motor neurons, as the “final common pathways” of the neuromuscular system (Sherrington (*16*), are the quanta of the neural control signals to muscles. In this study, we provided the first human dataset of the activity of dozens of motor neurons from six lower limb muscles during an isometric multi-joint task. We observed that only a small number of motor neurons was not significantly correlated to any of the other motor neurons (between 0 and 12%, which was participant-dependent). This proportion would likely be even lower if a more motor units could be identified. This result suggests that independent control of single motor units – if any – concerns an extremely small proportion of them. We contend that a pure independent control is unlikely, as it would be at the cost of a large computational capacity for the central nervous system, and would not be effective in regulating muscle force. Indeed, because the neural drive to the muscle corresponds to the sum of the discharge events of the activated motor neurons, the independent inputs received by the motor neurons are filtered out (*6, 11*). Therefore, any command signal should be common to a sufficient number of motor neurons to affect muscle force (*6*), and thus have a functional consequence.

### Grouping of motor neurons into functional clusters

The grouping of motor neurons into functional clusters supports the existence of a dimensionality reduction of the neural command at the spinal motor neuron level. This is in agreement with recent results obtained from intrinsic and extrinsic hand muscles (*4*). It forces a reconsideration of our understanding of the synergistic control of movement where the unit of control would be groups of motor neurons rather than muscles; and these groups of motor neurons and muscles may not necessarily overlap.

An important finding of this study is the partial mismatch between the distribution of common inputs to motor neurons (clusters) and muscle anatomy (Fig. 7). Although motor neurons from the same pool exhibited an overall high local density, we observed that some motor neurons from the same pool (i.e., innervating the same muscle) belonged to different clusters. This finding aligns particularly well with the observation that muscles may be spatially organized within discrete neuromuscular compartments (*17*). Each motor unit may generate a force having a particular direction, and therefore the recruitment of specific groups of motor units may be an effective strategy to comply with the mechanical constraints imposed by a given task. Some motor neurons from the same pool belong to different clusters, which can explain previous findings of an independent control of motor units from the same pool, leading to the assumption that motor neurons are flexibly controlled (*5*). We demonstrated that although some motor neurons from the same pool may receive different inputs, and therefore be controlled independently of each other, they would necessarily share common inputs with some other motor neurons, possibly from other pools.

We also identified common inputs to motor neurons from distant muscles, which aligns with our results obtained using bipolar EMG during dynamic pedaling and showing a synergy composed by the hamstrings and gastrocnemii (Fig. 1). It further supports previous results obtained using the classical synergy approach, and reporting the presence of distant muscles within the same synergy [e.g. hamstrings and plantarflexors during cycling (*18*), as well as gluteii and quadriceps during gait (*14*)]. However, the limitation of the interference EMG to assess neural drive made it impossible to verify that these distant muscles actually shared neural inputs. Using an *in vivo* approach in humans, our work demonstrates that motor neurons from distant muscles may receive a common input. The abovementioned finding was mainly observed between motor neurons of hamstrings and plantarflexors, which aligns well with the combined role of these muscles in lower limb extension (*19*). Further, this is compatible with a hierarchical control of multi-joint motor tasks (*20*). Of note, synchronized activity of motor neurons has been previously observed in a rat spinal cord preparation from pools located several spinal segments apart (*21*). However, this synchronization was less frequently observed than that between motor neurons from the same spinal segment (*21*), which aligns well with our observations.

In our study, motor neurons from distant pools exhibited an overall lower local density than those from anatomically related muscles, such as ST-BF, GM-GL and VL-VM pairs (Fig. 6). Specifically, motor neurons from the lateral and medial heads of the quadriceps densely overlapped (Fig. 5), suggesting that these two muscles might be controlled as a single muscle, at least during the studied motor task. This is probably because these two muscles share the same function as knee extensors, coupled with an important combined role in internal joint stress regulation (*22*). Of note, the fact that VL and VM are always coactivated may favor neural connection, which in turn triggers a strong common input between their motor neurons.

### Neural substrate for a synergistic control at the motor neuron level

Our approach did not allow us to determine the origin of the common synaptic input; nonetheless, our results are compatible with the notion that the spinal cord contains premotor interneurons that project to multiple motor neuron pools, including distant pools (*23, 24*), known as the neuron’s “muscle field” (*25*). Specifically, the work by Levine et al. (*24*) highlighted that some spinal neurons can cross multiple spinal segments to link distal functional motor neuron pools. Further, Takei et al. (*26*) observed that muscle fields of premotor interneurons are not uniformly distributed across muscles but rather distributed as clusters corresponding to muscle synergies. However, a degree of flexibility may exist beyond structural connections (hardwired in anatomical circuits), such that networks of motor neuron inputs adapt to different functional demands.

### Methodological considerations

Some methodological aspects should be considered in our experiment. First, we did not test for the robustness of the networks across different motor tasks because it required to identify the same motor units across tasks. It is particularly challenging to track motor units across tasks with different mechanical constraints given that muscle activation level and muscle geometry may change as a result of change in joint angles / force direction. Even though future studies should tackle this challenge, we do not believe that the absence of other motor tasks affect our main conclusion that the dimensionality of control of this particular multi-joint task was reduced by distributing common inputs to groups of motor neurons. Furthermore, an important conclusion, which is not affected by the absence of another motor task, is that groups of motor neurons were partly decoupled from their innervated muscle. Second, we studied a purely force-matched isometric task, thereby ensuring that the correlation between the smoothed spike trains reflected neural strategies rather than co-activation of motor units dictated by the task constraints. Indeed, in the cases where the muscle force is cyclically modulated during anisometric or isometric tasks, correlation is biased toward the cyclic time-varying force modulation, and thus from mechanical constraints (i.e., the co-activation of two muscles with similar functions, rather than a shared common input). Therefore, our results should not be extrapolated to other motor tasks, although we believe that our study was a necessary first step to demonstrate the synergistic control of spinal motor neurons. Finally, it is tempting to make comparison between participants, but we should keep in mind that we identified a small proportion of the active motor neurons. Even though differences in both the functional networks and the cluster composition between participants (Fig. 5 and 7) aligns with previous work suggesting the existence of individual muscle activation signatures (*27, 28*), we are not confident in making interpretations with the relatively small number of motor neurons analyzed in our study.

## Conclusion

Our study supports the theory that movements are produced through the control of small numbers of groups of motor neurons via common inputs and that these groups do not necessarily overlap with the innervated muscles. In this view, a common input is an essential feature of the neural control at the motor neuron level. Flexible grouping of motor neurons by distribution of common inputs allows for a large dimensionality reduction with respect to the available number of motor neurons, as well as a large range of variability in motor tasks due the large number of potential groupings. This theory explains observations on dimensionality reduction at the muscle level (*1, 2*), as well as of flexible control of motor neurons within the same pool (*5*).

## Methods

The entire data set (raw and processed data) is available at: https://figshare.com/s/dc7ce2758e4f3bbe6795

### 1. Participants

Ten males participated in this study (mean±standard deviation; age: 31.6±6.8 years, height: 181±5 cm, and body mass: 76±8 kg). Notably, we were not avoiding recruiting females; but decomposition of motor units is often more challenging in females (*29*). Participants had no history of lower leg pain that had limited function that required time off work or physical activity, or a consultation with a health practitioner in the previous six months. The ethics committee ‘CPP Ile de France XI’ approved the study (approval number: CPP-MIP-013), and participants provided an informed written consent.

### 2. Experimental design

An experimental session involved an isometric multi-joint task, wherein participants were instructed to match a force vector with the right leg on an instrumented clipless pedal, while seated on a cyclo-ergometer (Excalibur Sport; Lode, Groningen, the Netherlands). For this task, the right crank was fixed at 135° from the top dead center (Fig. 1). This crank angle was chosen as it is associated with the combined action of hamstrings and gastrocnemii to produce a pedal force (*18*), thereby allowing us to test the hypothesis of the presence of a common input between these distant muscles. Using a real-time force feedback, the participants were instructed to produce a force vector identical to that produced during a previously recorded dynamic cycling task performed on the same cyclo-ergometer. Specifically, they pedaled at 175W at 50 rpm for 90 s. The total reaction force applied on the right pedal was averaged across 15 cycles and both the pedal angle relative to the crank and the force vector produced at 135° were provided as a feedback for the isometric task described above. Participants were instructed to match the horizontal and vertical components of the force with an accuracy of ±15 N and the pedal angle with an accuracy of ±5°. Participants performed two contractions, each lasting 40 s, interspaced by 10 s of rest. Feedback of the target force vector and actual force vector was displayed on a monitor (Fig. 1). The overall aim of this approach was to study an isometric multi-joint task, which reproduced, as much as possible, the muscle coordination strategies used during dynamic pedaling. Importantly, a low pedaling rate was adopted such that non-muscular components of the pedaling force were minimized.

After the completion of the protocol described above, HDsEMG electrodes were replaced with classical bipolar EMG electrodes (Trigno Flex; Delsys, Boston, MA) and the participants were instructed to pedal at 175W at 50 rpm for 90 s. Muscle synergies were extracted using non-negative matrix factorization as described in Hug et al. (*18*). Of note, this analysis was performed on nine participants.

### 3. Pedal force measurements

The instrumented pedal was equipped with a LOOK Keo clipless platform (VélUS group, Department of Mechanical Engineering, Sherbrooke University, Canada). Participants wore cycling shoes. The sagittal plane components of the total reaction force applied at the shoe/pedal interface were measured using a series of eight strain gauges located within each pedal. The total reaction force was calculated from the measured Cartesian components F_T_ and F_N_, corresponding respectively to the horizontal forward and vertical upward forces on the pedal, respectively. An optical encoder with a resolution of 0.4° was used to measure the pedal angle. The top dead center was detected through transistor-transistor logic rectangular pulses delivered at the highest position of the right pedal. Zero adjustments for both components of force and pedal angle were performed before each session. Mechanical signals were digitized at 1000 Hz (DT 9800, Data translation, USA) and low-pass filtered at 5 Hz (2^nd^ order Butterworth filter).

### 4. High-density surface electromyographic recordings

HDsEMG signals were recorded from six lower limb muscles of the right leg: Biceps femoris (long head, BF), Semitendinosus (ST), Gastrocnemius medialis (GM), Gastrocnemius lateralis (GL), Vastus lateralis (VL), and Vastus medialis (VM). A two-dimensional adhesive grid of 64 electrodes (13×5 electrodes with one electrode absent on a corner, gold-coated, inter-electrode distance: 8 mm; [GR08MM1305, OT Bioelettronica, Italy]) was placed over each muscle. B-mode ultrasound (Aixplorer, Supersonic Imagine, France) was used to locate the muscle borders. Before electrode application, the skin was shaved, and then cleaned with an abrasive pad and alcohol. The adhesive grids were held on the skin using semi-disposable bi-adhesive foam layers (SpesMedica, Battipaglia, Italy). Skin-electrode contact was assured by filling the cavities of the adhesive layers with conductive paste. A 10-cm-wide elastic band was placed over the electrodes with a slight tension to ensure that all the electrodes remained in contact with the skin throughout the experiment. Strap electrodes dampened with water were placed around the contralateral (ground electrode) and ipsilateral (reference electrode for the gastrocnemius muscles) ankles. A reference electrode for the thigh muscles (5 × 5 cm, Kendall Medi-Trace™, Canada) was positioned over the patella of the right limb. The EMG signals were recorded in monopolar mode, bandpass filtered (10-500 Hz) and digitized at a sampling rate of 2048 Hz using a multichannel acquisition system (EMG-Quattrocento; 400-channel EMG amplifier, OT Biolelettronica, Italy).

### 5. Data analysis

#### HDsEMG decomposition

First, the monopolar EMG signals were bandpass filtered between 20-500 Hz with a second-order Butterworth filter. After visual inspection, channels with low signal-to-noise ratio or artifacts were discarded. The HDsEMG signals were then decomposed into motor unit spike trains using convolutive blind-source separation as previously described (*30*). In short, the EMG signals were first extended and whitened. Thereafter, a fixed-point algorithm that maximized the sparsity was applied to identify the sources of the EMG signals, i.e., the motor unit spike trains. The spikes were separated from the noise using K-mean classification and a second algorithm refined the estimation of the discharge times by minimizing the coefficient of variation of the inter-spike intervals. This decomposition procedure has been validated using experimental and simulated signals (*30*). After the automatic identification of the motor units, duplicates were removed and all the motor unit spike trains were visually checked for false positives and false negatives (*29, 31*). This manual step is highly reliable across operators (*32*). As classically done, only the motor units which exhibited a pulse-to-noise ratio > 30 dB were retained for further analysis (*33*). This threshold ensured a sensitivity higher than 90% and a false-alarm rate lower than 2% (*34*).

#### Correlation between smoothed motor unit spike trains

We estimated the correlation between motor unit smoothed discharge rates to assess the common inputs to motor neurons. To this end, the decomposed motor unit spike times were first converted into continuous binary signals with “ones” corresponding to the firing instances of a unit (Fig. 2A). The smoothed discharge rates were then obtained by convoluting these binary signals with a 400-ms Hanning window. Finally, these signals were high-pass filtered with a cut-off frequency of 0.75 Hz to remove offsets and trends (Fig 2B) (*35*). The level of common input for each pair of motor unit was estimated using a cross-correlation function applied on the smoothed discharge rates. The maximum cross-correlation coefficients within a time lag of −250 to +250 ms were considered and gathered to generate a 2-D correlation matrix (Fig. 2C).

The main analysis was performed on a 10-s window that was chosen such that the number of active motor units per muscle was maximized. We also assessed the repeatability of this analysis by comparing two 10-s windows selected during the first and second contractions. These two windows were chosen such that they contained the same units and that the number of active motor units per muscle was maximized. Because of the constraints of matching the motor units between the two contractions, the total number of motor units in this analysis was lower than that in the main analysis.

#### Networks of inputs to motor neurons

To derive the motor neuron networks, we used a threshold on the strength of the correlation between motor unit activities. This threshold was defined as the 99^th^ percentile of the cross-correlation coefficient distribution generated with resampled versions of the motor unit spike trains. Specifically, for each motor unit we generated a surrogate spike train by bootstrapping (random sampling with replacement) the interspike intervals. This random spike train had the same number of spikes, and the same discharge rate (mean and standard deviation) as the original motor unit spike train. Two iterations of this random procedure were performed, such that each motor unit was associated to two surrogate spike trains and each motor unit pair to four combinations of surrogate spike trains, thereby leading to 25,553 correlation coefficients for the whole population. This analysis was performed on the same 10-s window as that used for the main analysis, and led to a significant threshold of 0.35.

Similar to what has been proposed by Boonstra et al. (*36*) at the level of individual muscles, we used graph theory to identify motor neuron networks. Specifically, after determining the threshold, the Fruchterman-Reingold algorithm implemented in Origin Pro (2021b, OriginLab Corporation, Northampton, MA, USA) was used to allocate each motor neuron in a two-dimensional space (*12*). This method builds a graph based on the connectivity between nodes (in our case, a correlated activity of motor neurons) and on optimization criteria, such as even node distribution and uniform edge length. Each edge connecting nodes is considered as a spring with attractive and repulsive forces depending on its length, and the algorithm optimizes node positions to minimize the total energy of the network.

The generation of the graph according to the above-described procedure does not include any *a priori* information on motor neuron grouping (e.g., innervated muscles) and is solely based on the significant correlations of motor neuron activities. Therefore, the procedure is purely signal-based without any physiological or anatomical constraint.

#### Hierarchical clustering

After building the graphs of common inputs to motor neurons, we applied a clustering procedure to group the motor neurons based on the common input they received, as proposed at the level of individual muscles (*37*). A cluster was defined as a group of motor neurons densely connected to each other and loosely connected to the rest of the network. Each cluster was therefore a group of motor neurons with a high number of significant correlations between their activities and a minimum number of significant correlations with the activities of other motor neurons outside the cluster. The metrics that quantifies the strength of division of a network into multiple clusters is the modularity. Therefore, many algorithms that aim at identifying clusters within a network randomly assign nodes to clusters to iteratively maximize the modularity [e.g., the Louvain algorithm, (*38*)]. However, a well-known limitation of this approach is that it is stochastic, with numerous alternative results depending on the number of iterations or the arbitrary selection of a resolution parameter to estimate the modularity (*39*). To overcome this limitation, we used the multiresolution consensus clustering approach (*40*). First, we generated 1,000 partitions that cover the entire range of resolutions, i.e., from a partition where all the motor neurons belong to the same cluster to a partition where the number of clusters is equal to the number of motor neurons. This approach ensures an approximately equal coverage of all scales of the network. Second, we applied consensus clustering on the entire set of partitions (*41*). This step is an iterative procedure that consists of: 1) estimating the probability for each pair of motor neurons to belong to the same cluster across the partitions, resulting in a consensus matrix; 2) identifying clusters within this consensus matrix with a graph-clustering algorithm that optimizes the modularity, i.e., the GenLouvain algorithm (*42*), to generate a new set of partitions; and 3) repeating 1) and 2) until the procedure converges toward a unique partition (*41*). This “consensus partition” is considered as the most representative of all the partitions. It is also noteworthy that this algorithm only considers statistically significant consensus partitions. This means that the clusters of the ‘consensus partition’ cannot be identified in random networks generated by locally permutating the connections between motor neurons (*40*). Finally, we applied consensus clustering to each set of motor neurons embedded in newly generated clusters. This final step enabled us to identify clusters of motor neurons at multiple scales (Fig.2).

#### Network measures

We first assessed whether the data-driven networks were repeatable when computed from different time intervals. We assessed the repeatability of each network with four metrics: the mean node degree, density, global clustering coefficient, and modularity. The mean node degree is the average number of edges per node. A high mean node degree value signifies that the motor neurons within the network tend to connect (i.e. to be correlated) to many other motor neurons. The density is the fraction of actual connections (correlation above the threshold) to possible connections (the total number of pairs of motor neurons). The global clustering coefficient assesses the probability that each node is connected to its neighbors. It depicts the tendency of motor neurons to form groups based on their degree of common input. The modularity describes the strength of division of a network into multiple clusters. A high modularity value indicates that the number of significant correlations between motor neurons within each cluster is much higher that the number of significant correlations between motor neurons from different clusters. Importantly, the modularity depends on the value of a resolution parameter that defines the size of the identified clusters. Here, we used a fixed resolution parameter of 0.5. We calculated the intraclass correlation coefficient (ICC) for each metric to measure the repeatability.

After we considered networks of motor neurons without any *a priori* on their innervation, we explored the correspondence between the network measures and muscle anatomy. Specifically, we calculated the local density for each muscle and each pair of muscles, referred to as “subset”. To this end, we calculated the ratio between the number of significant correlations within the subset of motor neurons and the total number of possible connections within the subset of motor neurons. A high local density indicates a high degree of common input within the subset of motor neurons (muscle or muscle pair). A one-way analysis of variance was performed on the local density to assess the differences between subset of motor neurons. Multiple comparisons were performed using the Bonferroni approach.

To describe the reduced dimensionality of motor control during the isometric tasks, we reported the number of clusters in the last consensus partition. As described above, this last partition represents the smallest scale to which no additional significant clusters can be added.

## Acknowledgements

François Hug is supported by a fellowship from the *Institut Universitaire de France* (IUF). Dario Farina received support from the European Research Council Synergy Grant NaturalBionicS (contract #810346).

## Author Contributions

Contribution and design of the experiment: FH, SA, AdV, DF Collection of data: FH, AS

Analysis and interpretation: FH, SA, AdV, DF

Drafting the article or revising it for important intellectual content: FH, SA, AdV, DF All authors approved the final version of the manuscript.

